# Holder Pasteurization Inactivates SARS-CoV-2 in Human Breast Milk

**DOI:** 10.1101/2020.06.17.155689

**Authors:** Carina Conzelmann, Rüdiger Groß, Toni Luise Meister, Daniel Todt, Adalbert Krawczyk, Ulf Dittmer, Steffen Stenger, Jan Münch, Eike Steinmann, Janis A Müller, Stephanie Pfaender

## Abstract

SARS-CoV-2 RNA has been detected in the human breast milk of infected mothers, raising concerns regarding the safety of breastfeeding upon infection. We here show that holder pasteurization inactivates SARS-CoV-2 and provides an alternative and safe option for infected mothers to continue feeding breast milk to their infants.

## Introduction

The current coronavirus pandemic, caused by the severe acute respiratory syndrome coronavirus 2 (SARS-CoV-2), raises unprecedented questions regarding virus transmission and risks for pregnant or breastfeeding women. We and others recently described that SARS-CoV-2 is detectable in breast milk of infected mothers ^1–5^ and found viral RNA in single or multiple breast milk samples of mothers suffering from coronavirus disease 2019 (COVID-19). In two cases where the mother continued breastfeeding, the newborns were also tested positive for SARS-CoV-2 and one infant developed severe respiratory disease ^1,5^. However, the origin of the infections of the newborns remained unclear and raises concerns of possible virus transmission via breast milk.

The safety and feasibility of breastfeeding is of high importance as breast milk contains nutrients, hormones, and immunoprotective entities that are essential for the development, health, and protection of the neonate from infections ^6^. So far, the World Health Organization (WHO) recommends to continue breastfeeding upon SARS-CoV-2 infection of the mother while taking measures of strict hygiene and wearing masks to protect the child from droplets or aerosols ^7^. The Centers for Disease Control and Prevention (CDC) states that the decision for breastfeeding lays with the mother, family and health care providers ^8^. Generally, stopping breastfeeding is not advised, however, given the recent detection of SARS-CoV-2 in breast milk samples, a possible transmission of the virus via breastfeeding cannot be ruled out. Therefore, measures to provide safety of infants are urgently required. Thus, we here explored the inactivation of SARS-CoV-2 in human milk by holder pasteurization (heating to 63°C for 30 minutes) to reduce the risk of a possible virus transmission while preserving many of milk’s beneficial properties.

## Results

To test if SARS-CoV-2 retains infectivity in human breast milk and to explore holder pasteurization as a possible inactivation method, we spiked five different SARS-CoV-2 isolates from Germany, France and the Netherlands into five individual breast milk samples, and incubated them for 30 minutes at room temperature or 63°C. Samples where then titrated on cells and residual infectivity determined as tissue culture infectious dose 50 (TCID_50_). All five tested SARS-CoV-2 isolates (UKEssen, Ulm/01, Ulm/02, France/IDF0372, Netherlands/01) remained infectious in the milk samples that were incubated for 30 minutes at room temperature, with infectious titers of 0.09 to 1.2 × 10^5^ TCID_50_/mL (**Figure 1A, B**). Of note, in each milk sample, we detected a 40.9 – 92.8% decrease of viral titers compared to the medium control. This was independent of the viral strain, suggesting that this partial inactivation of virus is an intrinsic property of human milk, as previously described for many enveloped viruses including hepatitis C and Zika virus ^9–11^. Importantly, upon pasteurization, no residual infectivity was detected in any of the samples (**Figure 1A, B**). Thus, human breast milk containing infectious SARS-CoV-2 can be efficiently inactivated using standard holder pasteurization.

**Figure 1.**
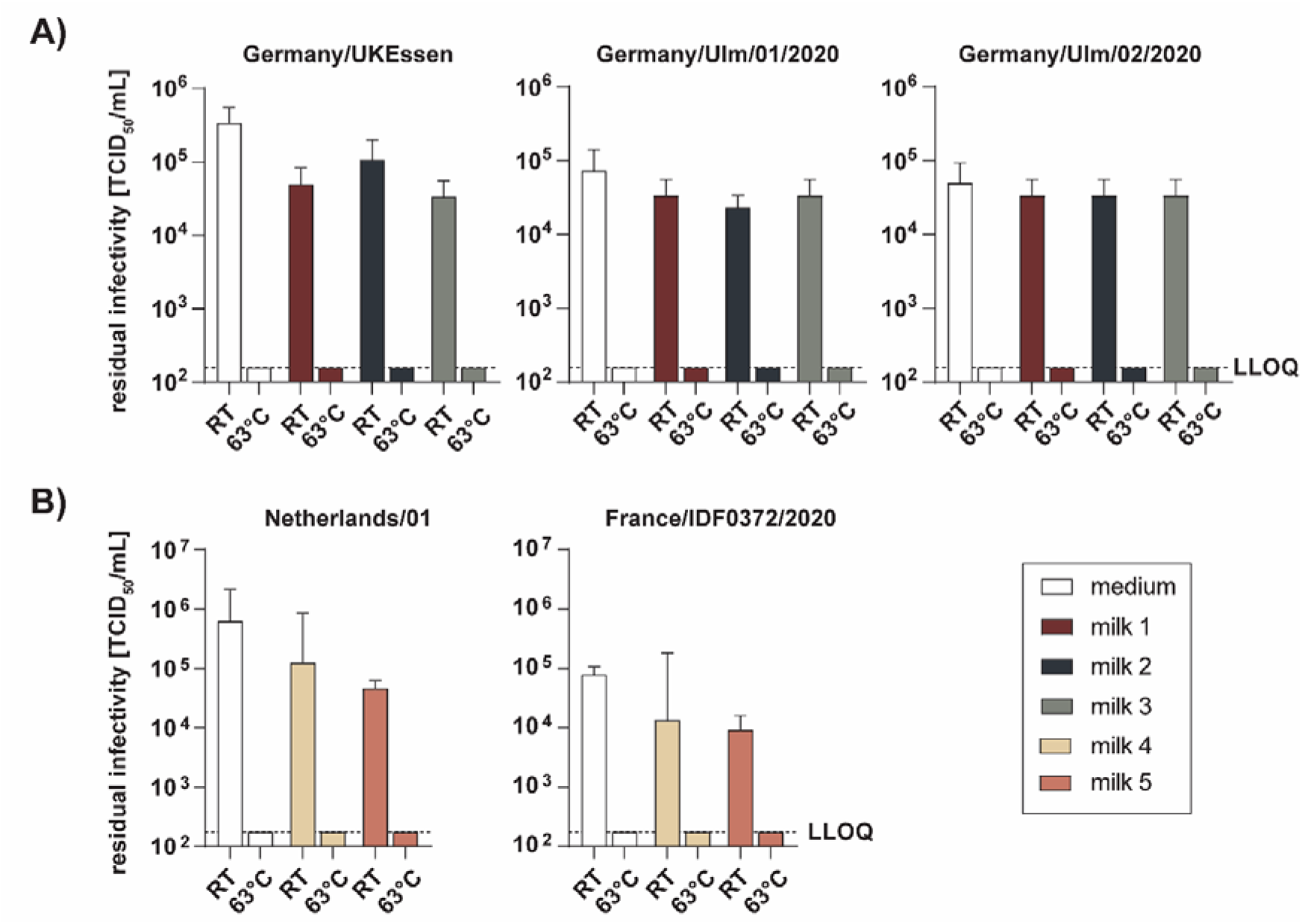
Holder Pasteurization inactivates SARS-CoV-2 in human breast milk. SARS-CoV-2 isolates UKEssen, Ulm/01, or Ulm/02 were spiked into medium or individual breast milk samples from donors 1-3 (**A**), or isolates France/IDF0372 and Netherlands/01 into milk samples from donors 4-5 (**B**), incubated for 30 minutes at room temperature (RT) or 63°C and titrated onto Vero E6 (**A**) or Caco-2 (**B**) cells to determine infectious titers. Tissue culture infectious dose 50 (TCID_50_) was calculated according to Spearman-Karber. Data indicate averages and standard deviation from two (B) or three (A) experiments. LLOQ lower limit of quantitation (A, 158 TCID_50_/mL; B, 176 TCID_50_/mL).

## Discussion

Breastfeeding provides many health benefits and is therefore recommended as the optimal feeding option for infants. Currently, the WHO recommends that SARS-CoV-2 infected mothers continue breastfeeding while taking measures of strict hygiene to protect the child from droplets or aerosols ^7^. As initial reports failed to detect the presence of viral RNA in milk samples of infected mothers ^12,13^, the risk of transmission via breastfeeding was deemed to be very low. Recent reports, however, demonstrate that SARS-CoV-2 RNA can be detected in breast milk samples after virus infection, associated with at least one newborn with severe respiratory disease ^1–5^. These findings raise the concern of safety during breastfeeding and a possible alternative or additional way of virus transmission to the infant. Holder pasteurization (heating to 63°C for 30 minutes) has been used for a long time to inactivate viral and bacterial agents, while at the same time preserving the many beneficial and protective effects of human breast milk ^6^. Here, we evaluated holder pasteurization as an easy and inexpensive methods to inactivate infectious SARS-CoV-2 in breast milk. Our data show that independent of the tested SARS-CoV-2 isolates or the breast milk sample, viral infectivity is completely eliminated by this treatment. Thus, holder pasteurization provides safety for the infant and reassurance for the mother, who might consider discontinuing breastfeeding and substituting for infant formula milk that lacks many of the human milk’s important components. In conclusion, we here provide a safe and feasible option, when in doubt, to continue feeding of the infant with breast milk upon symptomatic SARS-CoV-2 infection of the mother.

## Methods

### Cell culture

Vero E6 (*Cercopithecus aethiops* derived epithelial kidney) cells were grown in Dulbecco’s modified Eagle’s medium (DMEM, Gibco) which was supplemented with 2.5% heat-inactivated fetal calf serum (FCS), 100 units/mL penicillin, 100 μg/mL streptomycin, 2 mM L-glutamine, 1 mM sodium pyruvate, and 1x non-essential amino acids (DMEM complete). Caco-2 (human epithelial colorectal adenocarcinoma) cells were grown in DMEM complete but with supplementation of 10% FCS. All cells were grown at 37°C in a 5% CO_2_ humidified incubator.

### Virus strains and virus propagation

Viral isolate BetaCoV/France/IDF0372/2020 and BetaCoV/Netherlands/01 were obtained through the European Virus Archive global. The viral isolates BetaCoV/Germany/Ulm/01/2020, BetaCoV/Germany/Ulm/02/2020 and UKEssen were obtained from patient samples. Virus was propagated by inoculation of 70% confluent Vero E6 cells in 75 cm^2^ cell culture flasks with 100 μl SARS-CoV-2 isolates in 3.5 ml serum-free medium containing 1 μg/mL trypsin. Cells were incubated for 2 h at 37°C, before adding 20 ml medium containing 15 mM HEPES. Supernatant was harvested at day 3 post inoculation when a strong cytopathic effect (CPE) was visible. Supernatants were centrifuged for 5 min at 1,000 × g to remove cellular debris, and then aliquoted and stored at −80°C as virus stocks. Infectious virus titer was determined as plaque forming units or TCID_50_/mL.

### Milk samples

Milk samples were obtained from five healthy human donors after ethical approval by the ethics commission of Hanover Medical School, Hanover, Germany and the ethics committee of Ulm University. All mothers provided written informed consent for the collection of samples and subsequent analysis. Milk was collected freshly and stored at −80°C until further use as anonymized samples.

### TCID_50_ endpoint titration

To determine the tissue culture infectious dose 50 (TCID_50_), virus stocks or samples were serially diluted and used to inoculate Vero E6 or Caco-2 cells. To this end, 20,000 cells were seeded per well in 96 flat bottom well plates and incubated over night. Cells were infected with SARS-CoV-2 in serial dilutions and incubated for 3-6 days and monitored for CPE. TCID_50_/mL was calculated according to Spearman-Karber.

## Acknowledgements

We would like to thank all mothers and all members of the mid wife team Luna who contributed to the milk donation. Furthermore, we are grateful to all members of the Department of Molecular and Medical Virology of the RUB for the helpful discussions. We thank Lisa Schwefele MD (Div. of Neonatology, Dept. of Pediatrics, Ulm University Medical Center, Germany) and Frank Reister MD (Div. of Obstetrics, Dept. of Obstetrics & Gynecology, Ulm University Medical Center, Germany) for reviewing the manuscript from a clinical point of view. We thank Karin Steinhart MD (Administrative District Heidenheim, Public Health Office, Heidenheim, Germany) for organizing milk samples and Tirza Braun for experimental assistance. This project has received funding from the European Union’s Horizon 2020 research and innovation programme under grant agreement No 101003555 (Fight-nCoV) to J.M., the German Research Foundation (CRC1279) to J.M. and S.S., and an individual research grant (to J.A.M.). J.A.M. is indebted to the Baden-Württemberg Stiftung for the financial support of this research project by the Eliteprogramme for Postdocs. C.C., and R.G. are part of and R.G. is funded by a scholarship from the International Graduate School in Molecular Medicine Ulm. AK received funding from the Stiftung Universitätsmedizin Essen and the Rudolf Ackermann Foundation.

